# Keratin 17 regulates nuclear morphology and chromatin organization

**DOI:** 10.1101/2020.07.16.206730

**Authors:** Justin T. Jacob, Raji R. Nair, Brian G. Poll, Christopher M. Pineda, Ryan P. Hobbs, Michael J. Matunis, Pierre A. Coulombe

## Abstract

Keratin 17 (*KRT17*; K17), a non-lamin intermediate filament protein, was recently found to occur in the nucleus. We report here on K17-dependent differences in nuclear morphology, chromatin organization, and cell proliferation. Human tumor keratinocyte cell lines lacking K17 exhibit flatter nuclei relative to normal. Re-expression of wildtype K17, but not a mutant form lacking an intact nuclear localization signal (NLS), rescues nuclear morphology in *KRT17* null cells. Analyses of primary cultures of skin keratinocytes from a mouse strain expressing K17 with a mutated NLS corroborated these findings. Proteomics screens identified K17-interacting nuclear proteins with known roles in gene expression, chromatin organization, and RNA processing. Key histone modifications and LAP2β localization within the nucleus are altered in the absence of K17, correlating with decreased cell proliferation and suppression of GLI1 target genes. Nuclear K17 thus impacts nuclear morphology with an associated impact on chromatin organization, gene expression, and proliferation in epithelial cells.

**Summary:** Keratin 17 (K17) is one of two non-lamin intermediate filament proteins recently identified to localize to and function in the cell nucleus. K17 is here shown to regulate nuclear morphology, chromatin organization, LAP2 localization, and cell proliferation.

## Introduction

Intermediate filament (IF) proteins are encoded by a superfamily of >70 genes and assemble into fibrous cytoskeletal elements that fulfill key roles towards cellular structure and mechanics, and all other major cellular processes (Pan et al., 2013, Bouameur and Magin, 2017). Keratins, the largest subset of IFs, comprise 28 type I (acidic) and 26 type II (basic) keratin genes and proteins that form IFs as obligate heteropolymers in the cytoplasm of epithelial cells (Schweizer et al., 2006, Jacob et al., 2018). We and others recently discovered that select keratins and other non-lamin IF proteins can localize to the nucleus, contain functional importin α-dependent NLS sequences, and participate in important nuclear processes such as transcriptional control of gene expression, cell cycle progression, and protection from senescence (Escobar-Hoyos et al., 2015, Hobbs et al., 2015, Hobbs et al., 2016, Kumeta et al., 2013, Zhang et al., 2018, Zieman et al., 2019).

Before these findings the generally accepted notion was that, with the notable exception of type V nuclear lamins, IF proteins were defined to strictly reside in the cytoplasm (Omary et al., 2004, Hobbs et al., 2016). Similar views were once held for other cytoskeletal proteins that have since been found to occur and function in the nucleus (Pereira et al., 1998, Yeh et al., 2004, Bettinger et al., 2004, Pederson, 2008, Akoumianaki et al., 2009, Castano et al., 2010). For example, actin adopts novel conformations in the nucleus where it participates in chromatin remodeling, transcription via association with all three RNA polymerases, binding to nascent mRNA transcripts, and mRNA export (Bettinger et al., 2004, Schoenenberger et al., 2005, Visa and Percipalle, 2010, Kapoor et al., 2013). There is indirect evidence suggesting that the ancestral IF protein was a nuclear lamin-like protein from which all modern-day cytoplasmic IF proteins descended from (Erber et al., 1998). Accordingly, it could be that modern-day keratin proteins, in addition to other non-lamin IF proteins, are performing some of the “ancient” roles associated with IF proteins. This is not an unheard-of concept given the case of nuclear actin (Kapoor et al., 2013).

Kumeta et al. (2013) conducted an unbiased screen to identify the molecular components of nuclear scaffolds and reported on the occurrence of several keratins in the nucleus (e.g. GFP-tagged keratins 7, 8, 17, and 18; also, endogenous K8), with most appearing as a soluble, non-filamentous haze. We showed that keratin 17 (K17) and the transcriptional regulator AIRE co-localize in the nucleus and physically associate with similar and specific NF-κB (e.g. p65 subunit) consensus binding sequence-containing segments of the promoter regions of tumor-relevant genes associated with inflammation (Hobbs et al., 2015). Escobar-Hoyos et al. (2015) reported that K17 promotes cell cycle progression by binding to a G_0_/G_1_-to-S-phase transition inhibitor (e.g. p27^KIP1^) in the nucleus and facilitating its nuclear export, after which it is degraded by the ubiquitin-proteasome system in the cytoplasm. We have since demonstrated that another keratin protein, K16, localizes to the nucleus of epithelial cells (Zieman et al., 2019). The type IV IF protein nestin also localizes to the nucleus where it binds to and stabilizes lamin A/C, thereby protecting cells from senescence (Zhang et al., 2018). In the case of K17 and nestin, functional importin α-dependent bipartite NLS sequences have been identified (Escobar-Hoyos et al., 2015, Hobbs et al., 2015, Hobbs et al., 2016, Zhang et al., 2018). Despite this progress, we are still in the early stages of understanding the function(s) and significance of non-lamin IF proteins in the nucleus. Here, we provide evidence that nuclear-localized K17 regulates nuclear morphology and chromatin organization.

## Results and Discussion

### Loss of K17 alters nuclear morphology

To build on our knowledge of the presence, regulation, and function(s) of nuclear K17, we utilized CRISPR/Cas9 genomic editing technology to generate *KRT17* knock-out (KO) clones in two mammalian tumor-derived cell lines (HeLa, A431; see Methods). We observed a correlation between K17 expression and nuclear size and shape. From 2D-immunofluorescence images, loss of K17 correlated with larger nuclear area in HeLa (**Fig. S1 A,B**) and A431 (**Fig. S1 C**) human cells and in mouse epidermal keratinocytes (MEKs) in primary culture (**Fig. S1 D**). From confocal imaging, we observed that fewer optical sections were required to span the height of the nucleus (z-axis) in *KRT17* KO relative to parental cells (**Fig. 1 A**). Additionally, the nuclei of *KRT17* KO HeLa (**Fig. 1 B**), A431 (**Fig. 1 C**), and MEK (**Fig. 1 D**) cells exhibited decreased sphericity and increased total surface area, but no change in volume (**Fig. 1 B**).

**Figure 1.**
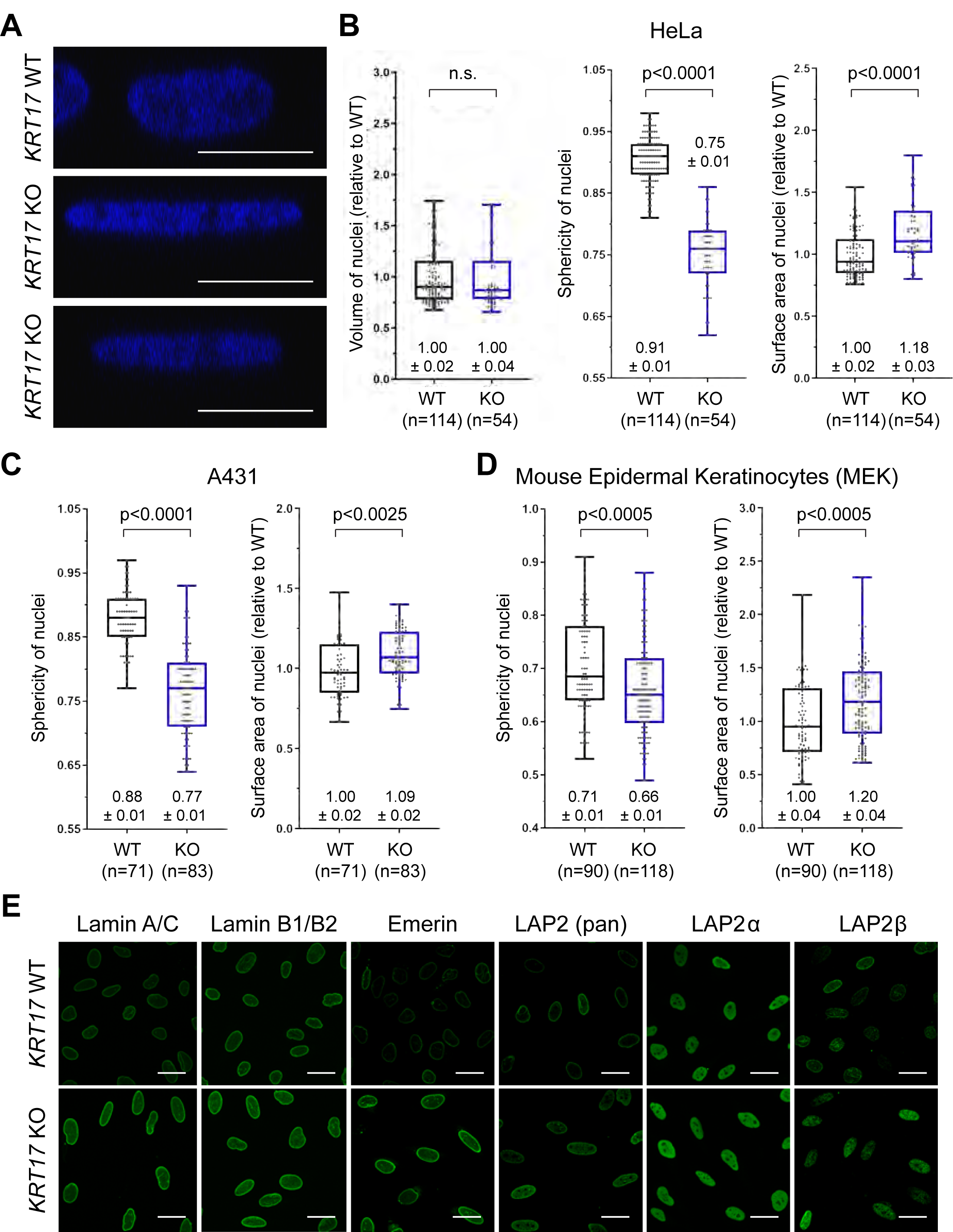
Loss of K17 alters nuclear morphology. **(A)** Orthogonal-view reconstruction of multi-plane confocal micrographs of individual nuclei from HeLa *KRT17* WT vs. *KRT17* knock-out (KO) cells. Nuclei are stained with DAPI (blue). Scale bars, 10 μm. **(B, C, D)** Quantitation of nuclear volume, sphericity, and /or surface area in (B) *KRT17* WT vs. KO cells HeLa and (C) A431 cells, and in (D) primary MEK cells. Numeric values above or below each box-and-whisker plot designate the mean +/- standard error of the mean (s.e.m.). **(E)** Confocal micrograph maximum intensity projections (MIPs) of nuclei from *KRT17* WT and *KRT17* KO HeLa cells, immunostained with antibodies recognizing several nuclear lamina-associated proteins (green). Scale bars, 10 μm.

We hypothesized that differences in nuclear shape and surface area may entail changes in inner nuclear membrane (INM) proteins, including lamin IFs (lamins A, C, and B1/B2) and LEM-domain-containing proteins (LAP2α, LAP2β, LAP2γ, emerin). Immunoblotting of whole-cell lysates from *KRT17* WT and KO cells revealed significant increases in the steady-state levels of all proteins analyzed, except LAP2β, in cells lacking K17 (**Fig. S1 E,F**). The observed increases (∼15-50%) were proportional to the increase in surface area (∼10-20%) in *KRT17* KO nuclei relative to control (**Fig. 1 B-D**). Consistent with these findings, indirect immunofluorescence microscopy indicated a significant increase in signal intensity for each analyzed protein in *KRT17* KO cells compared to WT (**Fig. 1 E**; quantitation in **Fig. S1 G)**. Again, LAP2β behaved differently (see below).

### Impact of K17 on nuclear morphology requires an intact NLS

We next assessed whether nuclear-localized K17 impacts nuclear dimensions. We transiently transfected *KRT17* KO cells (HeLa and A431) with either GFP (control), GFP-K17 WT or GFP-K17ΔNLS (in which K399A and K400A mutations impair NLS function; Escobar-Hoyos et al., 2015; Hobbs et al., 2015) (**Fig. 2 A**) and assessed the size and shape of nuclei from 2D and 3D immunofluorescence images. Expression of GFP-K17 WT, but not GFP-K17ΔNLS, restored normal dimensions to the nucleus, including decreased area (2-D; **Fig. 2 B,C**), increased sphericity (3-D; **Fig. S2 A,B**), and decreased surface area (3-D; **Fig. 2 D,E**). Parallel studies revealed that expression of GFP-K17 ΔNES in which L194A, L197A and L199A mutations impair K17’s NES function (thus favoring nuclear retention; Escobar-Hoyos et al., 2015) (**Fig. S2 C**) also rescues the nuclear shape defects occurring in *KRT17* KO A431 keratinocytes. Expression of GFP-K14 WT (**Fig. S2 D**) is ineffective in doing so, suggesting specificity for K17. Next, we used PLA assays with two different host species antibodies against K17 to relate the presence of K17 in the nucleus of HeLa *KRT17* KO keratinocytes transiently expressing GFP-K17 WT or GFP-K17ΔNLS. Both constructs gave rise to well-developed filamentous arrays in the cytoplasm of cells expressing them (see **Fig. 2 A**). As expected (Hobbs et al., 2015), GFP-K17ΔNLS gave rise to markedly reduced nuclear signal relative to GFP-K17 WT (**Fig. S2 E**). Taken together, these data suggest that, in transfected human A431 and HeLa cells, a nuclear pool of K17 is responsible for the impact on nuclear morphology, independently of the property of fostering a dense network of keratin filaments in the perinuclear cytoplasm.

**Figure 2.**
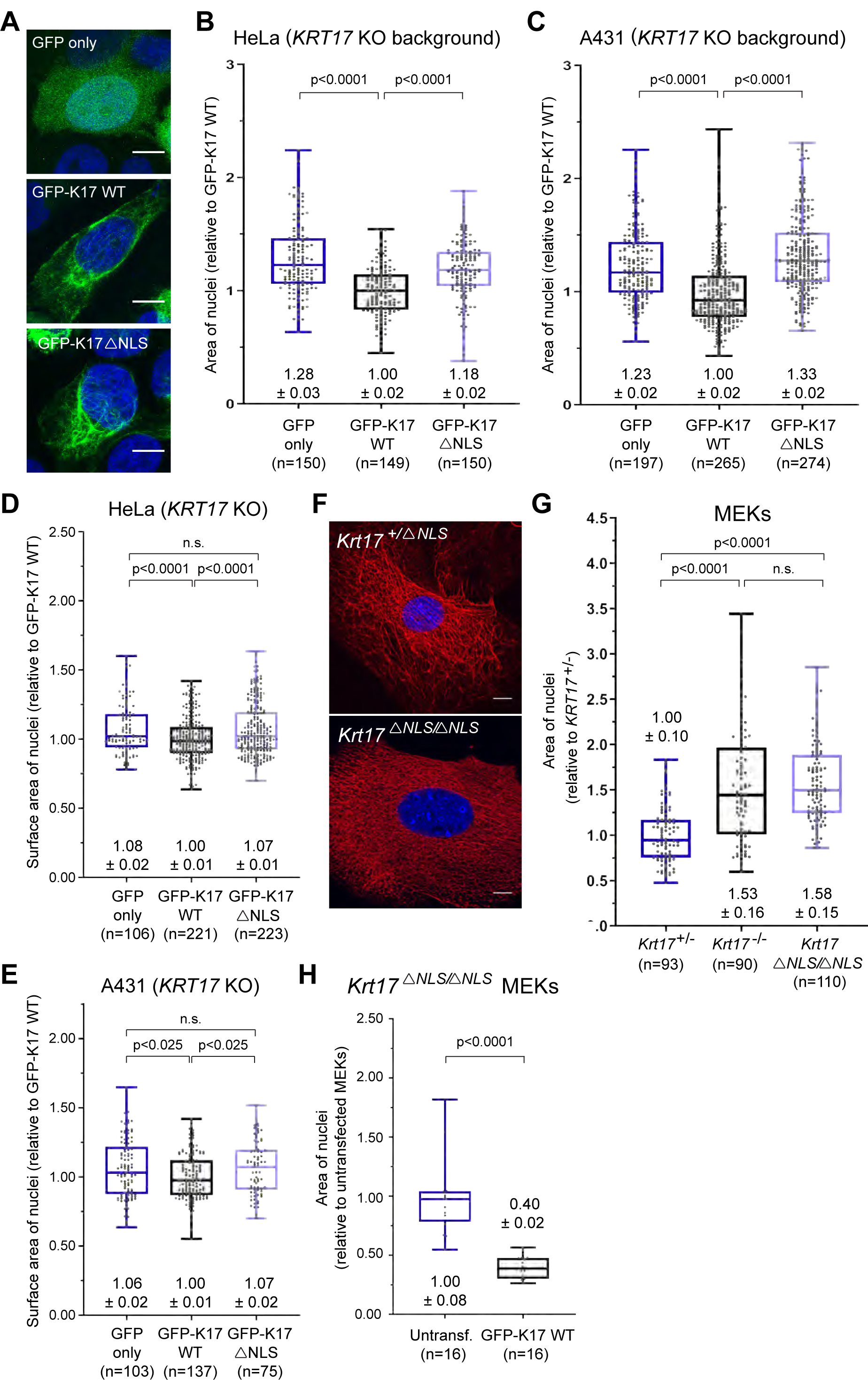
Impact of K17 on nuclear morphology requires an intact NLS. **(A)** Confocal micrograph maximum intensity projections (MIPs) of HeLa *KRT17* KO cells transiently transfected (48 h) with GFP (control), GFP-K17 WT, or GFP-K17ΔNLS plasmids (green). Nuclei are stained with DAPI (blue). Scale bars, 10 μm. **(B, C)** Quantified area measurements (2-dimensional; 2-D) of nuclei from (B) HeLa *KRT17* KO and (C) A431 *KRT17* KO cells transiently expressing GFP, GFP-K17 WT, or GFP-K17ΔNLS. **(D, E)** Quantified surface area (3-dimensional; 3-D) measurements of nuclei from (D) HeLa *KRT17* KO and (E) A431 *KRT17* KO cells transiently expressing GFP, GFP-K17 WT, or GFP-K17ΔNLS. **(F)** Confocal MIPs for K17 (red) in primary MEKs from *Krt17*^+/-^ and *Krt17*^*ΔNLS/ΔNLS*^ mice. Nuclei are stained with DAPI (blue). Scale bars, 10 μm. **(G)** Quantified area measurements of nuclei from primary MEKs from *Krt17*^+/-^, *Krt17*^-/-^, and *Krt17*^*ΔNLS/ΔNLS*^ mice. **(H)** Quantified area measurements of nuclei from *Krt17*^*ΔNLS/ΔNLS*^ MEKs either untransfected or transfected with GFP-K17 WT. For B, C, D, E, G and H, numbers above or below each box-and-whisker plot represent the mean +/- s.e.m.

We next analyzed nuclei in primary cultures of skin keratinocytes obtained from mice expressing a *Krt17*^*ΔNLS*^ allele. A CRISPR-Cas9-based strategy was used to produce a mouse strain carrying K399A and K401A mutations at the *Krt17* locus in its germline (**Fig. S3 A,B** and Methods). Mice homozygous for this allele (*Krt17*^*ΔNLS/ΔNLS*^) are viable, fertile, and show no obvious phenotype at baseline (**Fig. S3 C** and data not shown). The skin of young adult *Krt17*^*ΔNLS/ΔNLS*^ mice show normal histology relative to *WT* (**Fig. S3 D**) and normal pattern of immunostaining for K17 protein in hair follicles (**Fig. S3 E**). In primary culture, *Krt17*^*ΔNLS/ΔNLS*^ keratinocytes show a normal array of K17-containing filaments in the cytoplasm (**Fig. 2 F**). PLA assays (using two different host species antibodies for K17; allows the visualization of the K17 nuclear pool without LMB treatment) coupled to single-plane confocal microscopy show a markedly reduced frequency of nuclear-localized signal in *Krt17*^*ΔNLS/ΔNLS*^ mutant keratinocytes relative to wildtype (data not shown; similar to **Fig. S2 E**). Measurements revealed that, in spite of the presence of a robust network of K17 filaments in their cytoplasm (**Fig. 2 F**), the nuclei of newborn *Krt17*^*ΔNLS/ΔNLS*^ keratinocytes display a surface area that is indistinguishable from *Krt17* null keratinocytes but significantly larger than HET control (**Fig. 2 G**). Transfection-mediated re-expression of GFP-K17 WT, but not GFP-K17ΔNLS, restored nuclear dimensions in *Krt17*^*ΔNLS/ΔNLS*^ keratinocytes in primary culture (**Fig. 2 H**). Such findings extend our observations from cultured human cell lines to mouse skin keratinocytes in primary culture, and markedly strengthen the connection between nuclear-localized K17 and nuclear shape in skin keratinocytes.

### K17 interacts with proteins involved in chromatin organization and gene expression

To explore how K17 may regulate nuclear morphology, we conducted an unbiased screen to identify proteins that interact with K17 in the nucleus. We prepared a soluble subcellular fraction enriched for nuclear proteins, immunoprecipitated K17 from these lysates, and identified co-precipitated proteins by mass spectrometry (MS). Data filtering (e.g., 127N/130N ratio >1.2; 127N/130N count >4; see Methods) yielded 77 proteins that are enriched by K17 IP in *KRT17* WT cells over *KRT17* KO cells (see source data file). Of the 77 proteins, 58 (75%) had at least one defined nuclear role according to Gene Ontology terms in the Universal Protein Resource (UniProt) database (https://www.uniprot.org/). Network association analysis (STRING) under a high stringency setting (0.7) categorized these 77 MS-identified proteins as overrepresented in nuclear processes including RNA processing (n= 20), chromatin organization (n= 16), and regulation of gene expression (n= 41). RNA processing and chromatin organization each form a large hub in the network of functionally interconnected candidate nuclear protein interactors for K17 (**Fig. 3 A**).

**Figure 3.**
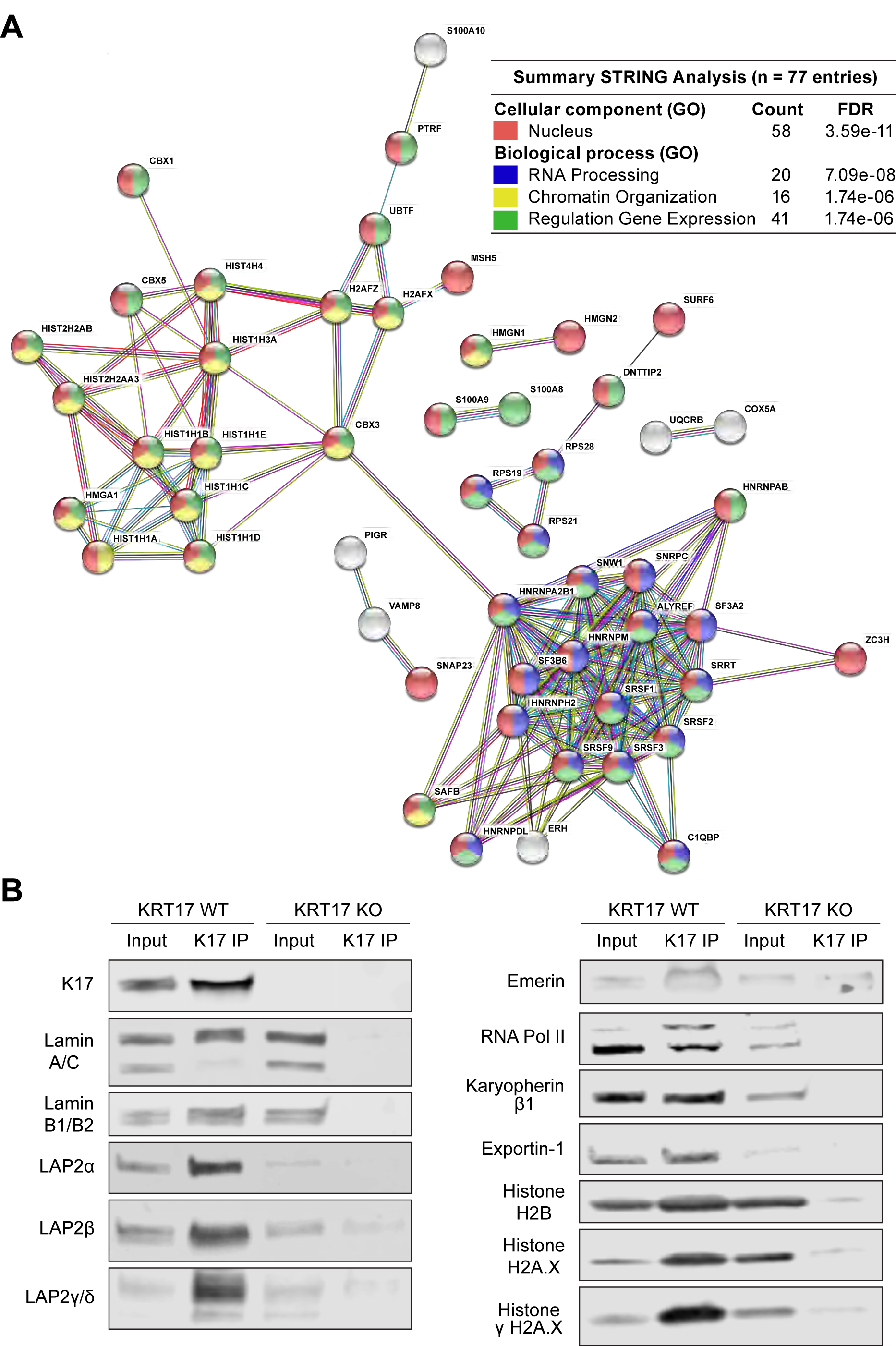
Screen for K17-interaction proteins in the nucleus. **(A)** STRING-db online analysis (Version 10.5; http://string-db.org/) of the 77 most-enriched MS-identified co-immunoprecipitated proteins, 58 (75%) of which have at least one known function in the nucleus, according to UniProt Gene Ontology (GO) Biological Process terms. Each circle or node represents one of the 77 candidate nuclear protein interactors, labeled with their gene names. Lines connecting two nodes designate a known functional protein-protein interaction, and line thickness indicates the strength of data supporting the interaction (note that only proteins with at least one connection are shown (n=52); this explains why K17 is not part of the network shown here. See source data file that accompanies this article). Red color reflects known nuclear localization; blue reflects a role in RNA processing; yellow reflects a role in chromatin organization, and green reflects a role in the regulation of gene expression (multi-coloring reflects more than one role). Inset table (upper right) highlights key findings from this analysis. “Count”, number of entries for a particular GO term (out of 77); “FDR”, False Discovery Rate. See text for details. **(B)** Western blots of several proteins (e.g. nuclear lamina-associated proteins, RNA polymerase II, nuclear import/export machinery, and histones) that co-immunoprecipitate with K17 in subcellular fractions enriched for nuclear proteins.

We next conducted a follow-up, target-specific survey that confirmed some of the MS results and identified several new K17-interacting proteins with well-defined roles in nuclear lamina architecture and chromatin organization. These included lamins A, B1, and B2; but not lamin C, which may indicate the potential for specificity. Also identified were LAP2 α, β, γ, and Δ isoforms; emerin; histones H2B, H2A.X, and γH2A.X; RNA Polymerase II; and the nucleo-cytoplasmic shuttling proteins exportin-1 and karyopherin β1 (**Fig. 3 B**). Similar results were obtained in HeLa and A431 keratinocytes.

### Loss of K17 alters histone modifications relevant to chromatin status and gene expression

Changes in nuclear size, shape, and/or lamina composition can lead to re-organization of chromatin-nuclear lamina interactions with an impact on gene expression (Mukherjee et al., 2016). Besides, the differential distribution of post-translational modifications (PTMs) on histone proteins are major determinants of chromatin and gene expression (Lawrence et al., 2016). We next tested whether histone PTMs are regulated by K17 expression status. Immunoblotting of whole-cell protein lysates from *KRT17* WT and KO HeLa cells revealed marked decreases in several specific histone PTMs, including acetyl-Histone H3 (at Lys-9 and/or Lys-14), acetyl-Histone H4 (at Ser-1, Lys-5, Lys-8, and/or Lys-12), and monomethyl-Histone H4 (at Lys-20), as a result of ablated K17 expression (**Fig. 4 A**; quantitation in **Fig. 4 B**). Confocal microscopy analyses, which afford an opportunity to assess individual cells rather than cell populations, confirmed reduced signal intensities of these histone PTMs in *KRT17* KO (human) cells relative to *KRT17* WT controls (**Fig. 4 C-F**). Altogether, K17 expression is correlated with more spherical nuclei and higher levels of histone PTMs associated with transcriptionally active euchromatin, which suggest that K17 may regulate gene expression epigenetically, e.g., through modifying the histone code.

**Figure 4.**
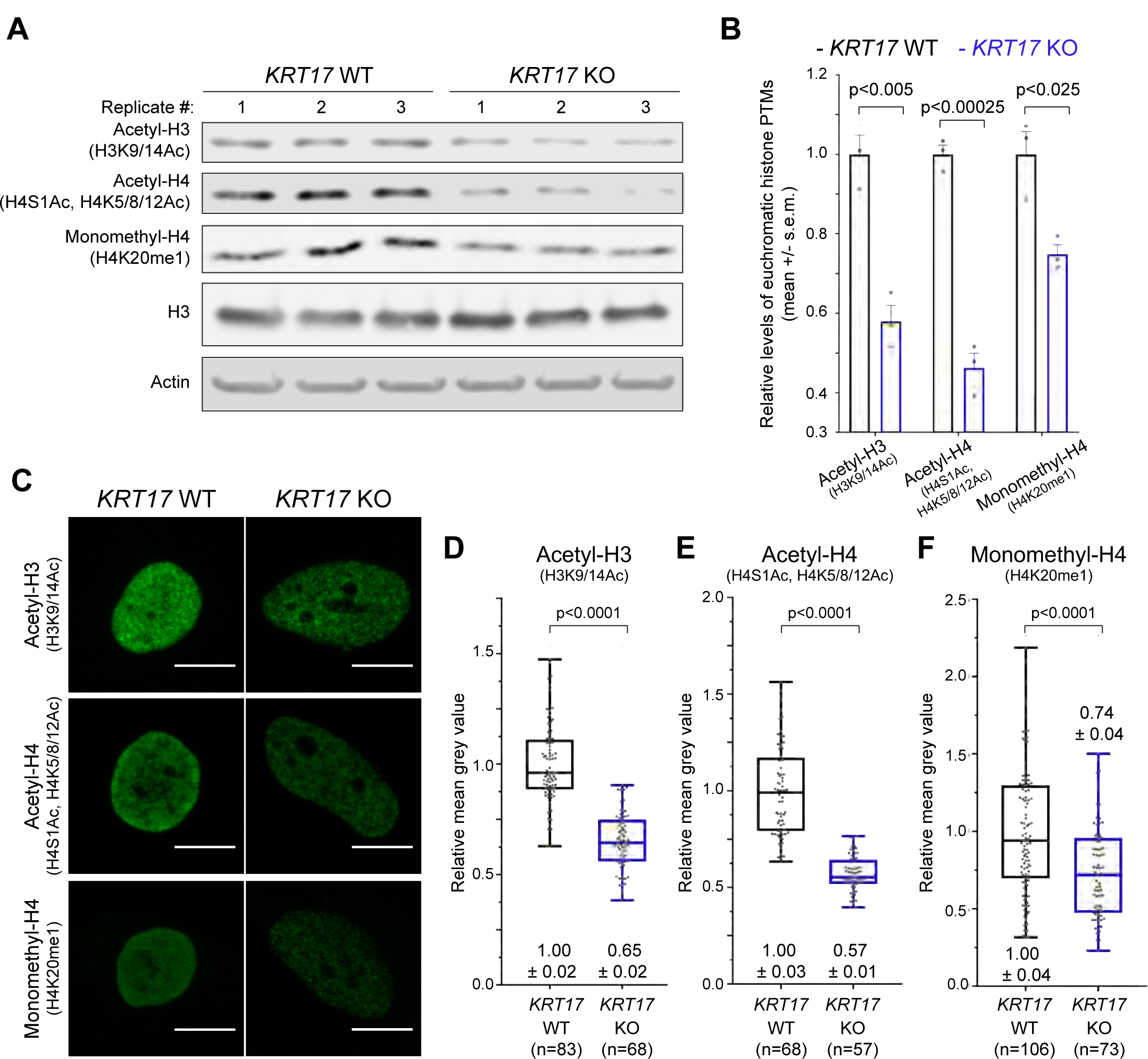
Loss of K17 affects histone modifications. **(A)** Western blots of whole-cell lysates from *KRT17* WT and *KRT17* KO cells (n=3 biological replicates per genotype). Primary antibody used is shown at left. **(B)** Quantitation of Western blot band intensities in (A), each normalized to β-Actin loading control. **(C)** Confocal micrograph maximum intensity projections of *KRT17* WT and *KRT17* KO cells immunostained with antibodies recognizing acetylated Histone H3 (H3K9/K14Ac), acetylated Histone H4 (H4S1Ac, H4K5/K8/K12Ac), or monomethylated Histone H4 (H4K20me1) (green). Scale bars, 10 μm. **(D, E**, and **F)** Quantitation of mean immunofluorescence signal intensity of (D) acetylated Histone H3 (H3K9/K14Ac), (E) acetylated Histone H4 (H4S1Ac, H4K5/K8/K12Ac), and (F) monomethylated Histone H4 (H4K20me1) in *KRT17* WT vs. *KRT17* KO cells. Numeric values above or below each box-and-whisker plot designate the mean +/- s.e.m. and arel relative to *KRT17* WT.

### Loss of K17 alters LAP2β localization, cell proliferation, and GLI1 function

LAP2β stood out by many criteria in our survey of nuclear envelope-associated proteins (see **Fig. 1 E** and **Fig. S1 E-G**). Follow-up quantitative confocal microscopy showed that LAP2β localization shifts from occurring in the INM and nucleoplasm of *KRT17*-positive cells to being predominantly in the nucleoplasm in *KRT17* KO cells (**Fig. 5 A,B**). A recent study showed that GLI1, a terminal effector of Hedgehog signaling and powerful mitogenic transcription factor (Mazza et al., 2013), is subject to a two-step, LAP2-dependent regulation when inside the nucleus (Mirza et al., 2019). After import, acetylated-GLI1 is initially docked to the INM in a LAP2β-dependent fashion and thus unable to drive expression of its target genes (Mirza et al., 2019), many of which drive cell proliferation (Mastrangelo and Milani, 2018). We surmised that the disruption of a potential interplay between K17, LAP2β, and GLI1 may explain previously reported differences in the proliferation properties of *Krt17* null keratinocytes (Hobbs et al., 2015, Depianto et al., 2010).

**Figure 5.**
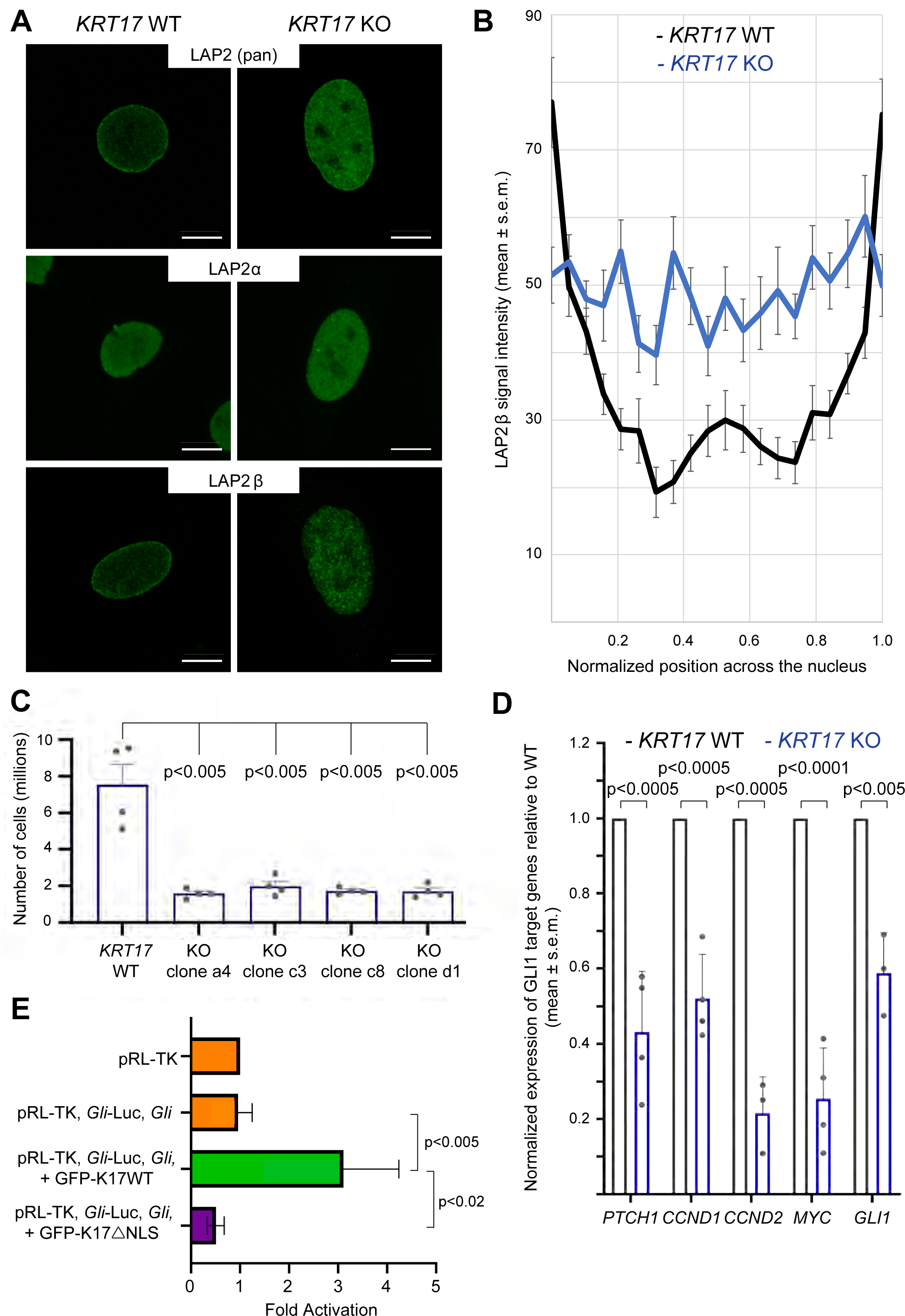
Loss of K17 and LAP2β, cell proliferation, and GLI1 function. **(A)** Confocal micrograph maximum intensity projections (MIPs) of *KRT17* WT and *KRT17* KO HeLa cells immunostained with antibodies recognizing nuclear lamina-associated proteins LAP2 (pan-isoform), LAP2α, and LAP2β (green). Scale bars, 10 μm. **(B)** Quantitation of signal intensity across line scans through individual nuclei from *KRT17* WT and *KRT17* KO HeLa cells (n=20 nuclei per genotype). Signal at the peripheries of the x-axis (i.e. normalized positions ∼0.00-0.05 and ∼0.95-1.00) reflect LAP2β signal at the nuclear envelope. Signal in the interior of the x-axis (i.e. normalized positions ∼0.05-0.95) reflect LAP2β signal in the nucleoplasm. **(C)** Cell counting assay using a hemocytometer to count the number of cells after seeding 500,000 cells per genotype (e.g. HeLa *KRT17* WT cells and four different HeLa *KRT17* KO clones) and culturing at 37°C and 5% CO_2_ for three days. **(D)** Normalized expression of select *GLI1*-target genes in *KRT17* KO relative to *KRT17* WT HeLa cells (n=3-4 biological replicates per mRNA target). **(E)** Luciferase assays in HeLa cells transfected with a *Gli1*-Luciferase reporter construct (see Methods). Data were normalized with regard to transfection efficiency and signal obtained with pRL-TK vector control. Data represent mean ± SEM from 5 biological replicates each consisting of 6 technical replicates. Mann-Whitney tests were performed to compare each parameter using GraphPad Prism 8. **P< 0.01, *P<0.05.

We examined proliferation in parental and *KRT17* null HeLa and A431 tumor cells. *KRT17* null HeLa and A431 cells proliferate more slowly relative to their respective parental controls, with the defect being more pronounced in HeLa (**Fig. 5 C**). To assess for cell-autonomy of this trait, we mixed and plated parental and *KRT17* null HeLa cells in a 1:1 ratio and monitored growth over time. Reproducibly, parental HeLa cells became dominant over time (**Fig. S3 F**). Further, use of the irreversible inhibitor of cell proliferation mitomycin C (Blake et al., 2006) showed that the two cell genotypes survived equally well (**Fig. S3 G**), ruling out that differential cell death contributed to the observed outcome. These findings are supported by data reported in DepMap, a survey of the impact of CRISPR-induced null alleles in a large collection of cell lines (https://depmap.org), showing that loss of K17 results in reduced proliferation in 324 out of 739 cell lines (note: loss of K14, a similar type I keratin, is neutral in this setting).

Next, we explored whether the proliferation deficit observed in *KRT17* null cells can be correlated to reduced GLI1 activity. First, we measured the mRNA levels for GLI1 target genes with an established role in cell proliferation. By RT-qPCR, the *PTCH1, CCND1, CCND2, MYC*, and *GLI1* mRNAs are each markedly decreased in *KRT17* null relative to parental HeLa cells (**Fig. 5 D**). Second, we conducted luciferase reporter assays using an established reporter of GLI1 transcriptional activity in *KRT17* KO HeLa cells (see Methods). Expression of GFP-K17 WT, but not GFP-K17ΔNLS, signficantly stimulated GLI1 reporter activity in *KRT17* KO HeLa cells (**Fig. 5 E**). These findings support the possibility that mis-localization of LAP2β in the nuclei of cells lacking K17 leads to reductions in GLI1-dependent gene expression and cell proliferation.

In conclusion, our findings establish a novel role for nuclear-localized K17 as a regulator of nuclear architecture, chromatin organization, and epigenetic modification of gene expression in keratinocytes. We report on >80 novel nuclear proteins (confirmed and candidates) in the “K17 interactome”, many of which have defined roles in chromatin dynamics, gene expression, RNA processing, nucleo-cytoplasmic trafficking, and DNA repair. These findings expand the knowledge of functions for nuclear keratins and points to exciting lines of future investigations.

## Acknowledgments

The authors are grateful to Amar Mirza, Anthony Oro, Beau Su, Daniela Drummond-Barbosa, Hoku West-Foyle, Loza Lee, Robert Cole, Robert Goldman, Roland Foisner, Scot C. Kuo, and Tatiana N. Boronina for their advice and support, and to Santa Cruz Biotechnology Inc. for providing antibodies. These studies were supported by grant R01AR44232 (to PAC) and grant T32GM007315 (to CMP) from the National Institutes of Health. The authors declare no competing financial interests.

## Supplementary Figures

**Figure S1** demonstrates that the loss of K17 alters the morphology of the cell nucleus and leads to increased levels of several nuclear lamina-associated proteins. **Figure S2** underscores the importance of K17’s NLS in re-establishing nuclear morphology. **Figure S3** highlights the generation, validation, and initial characterization of the *Krt17*^*ΔNLS/ΔNLS*^ genetic mouse model and co-culture experiments *with KRT17* WT and *KRT17* KO HeLa cells.

**Figure S1.**
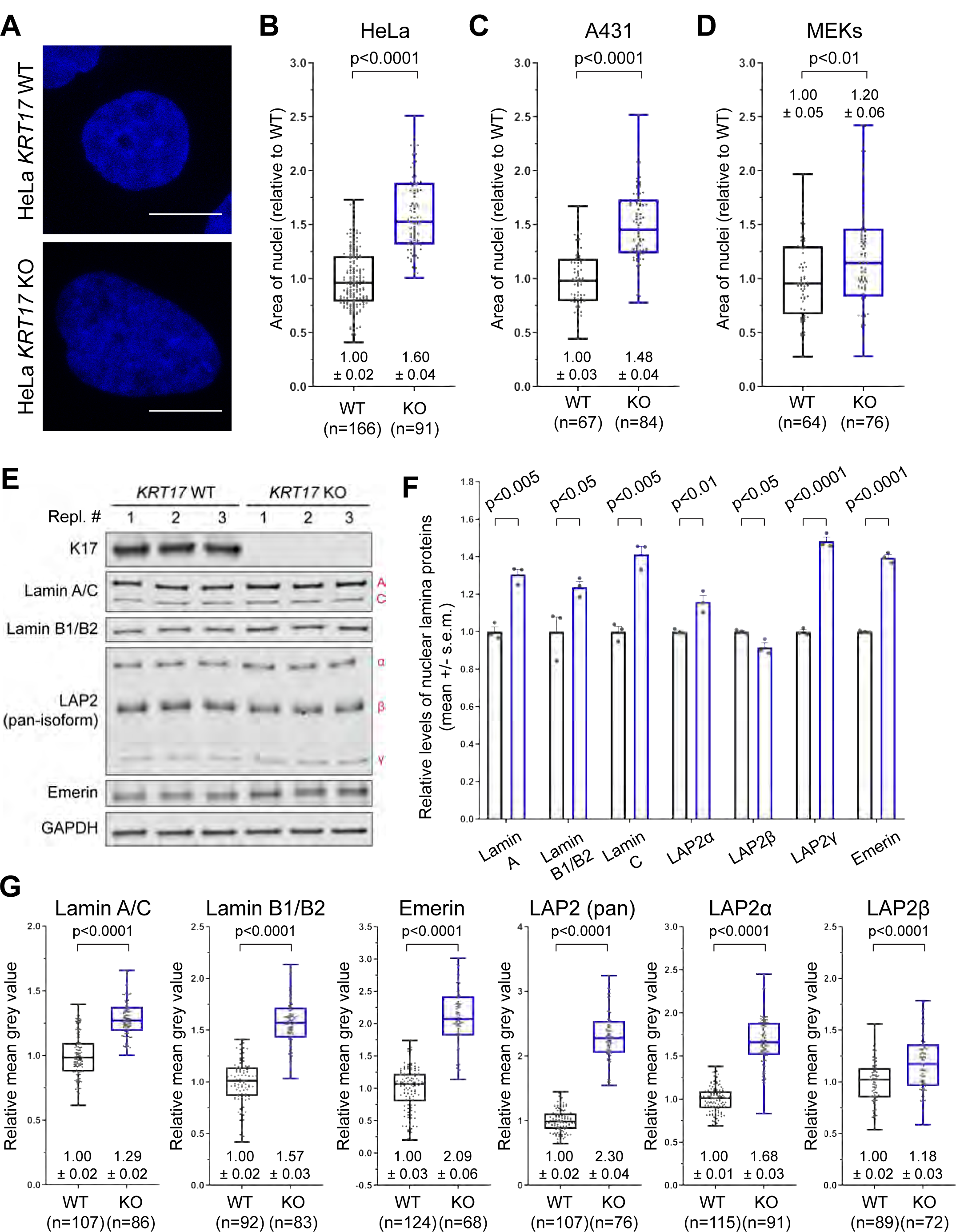
Loss of K17 alters nuclear morphology and nuclear lamina-associated proteins. **(A)** Confocal micrograph maximum intensity projections (MIPs) of individual nuclei from *KRT17* WT vs. *KRT17* KO HeLa cells. Nuclei are stained with DAPI (blue). Scale bars, 10 μm. **(B, C, and D)** Quantified area measurements of nuclei from cultured *KRT17* WT vs. KO *KRT17* (B) HeLa and (C) A431 human epithelial tumor cells, and (D) primary mouse epidermal keratinocytes (MEKs). Numeric values above or below each box-and-whisker plot designate the mean +/- standard error of the mean (s.e.m.). **(E)** Western blots of nuclear lamina-associated proteins from whole-cell lysates of *KRT17* WT and *KRT17* KO HeLa cells (n=3 biological replicates per genotype). **(F)** Quantitation of Western blot band intensities in (E), each normalized to GAPDH loading control. **(G)** Quantitation of mean immunofluorescence signal intensity of confocal micrograph MIPs of nuclei immunostained with antibodies recognizing several nuclear lamina-associated proteins (refer to **Fig. 1 E**).

**Figure S2.**
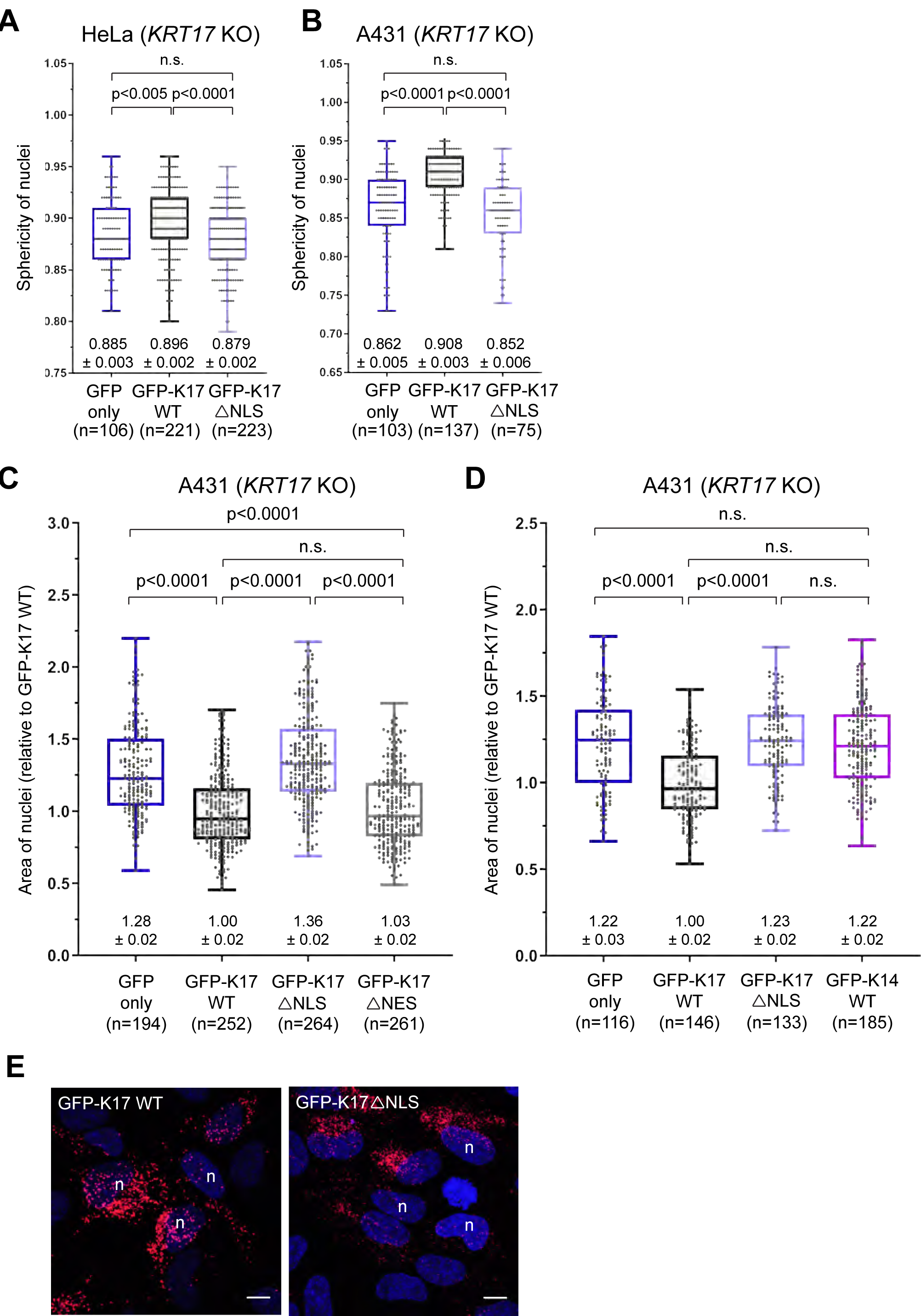
Nuclear-localized K17 rescues nuclear morphology. **(A and B)** Quantified sphericity measurements of nuclei from (A) HeLa *KRT17* KO and (B) A431 *KRT17* KO cells transiently transfected (48-hour transfection) with GFP (only), GFP-K17 WT, or GFP-K17ΔNLS plasmid constructs. Numeric values below each box-and-whisker plot designate the mean +/- s.e.m. **(C and D)** Quantified area measurements of nuclei from A431 *KRT17* KO cells transiently transfected (48-hour transfection) with GFP (only), GFP-K17 WT, GFP-K17ΔNLS, and (C) GFP-K17ΔNES or (D) GFP-K14 WT plasmid constructs. Numeric values below each box-and-whisker plot designate the mean +/- s.e.m. **(E)** Confocal micrograph maximum intensity projections of proximity ligation assays (PLAs) of HeLa *KRT17* KO cells transfected (48-hour transfection) with GFP-K17 WT or GFP-K17ΔNLS. PLAs relied on the use of two different host species antibodies recognizing K17. Scale bars, 10 μm.

**Figure S3.**
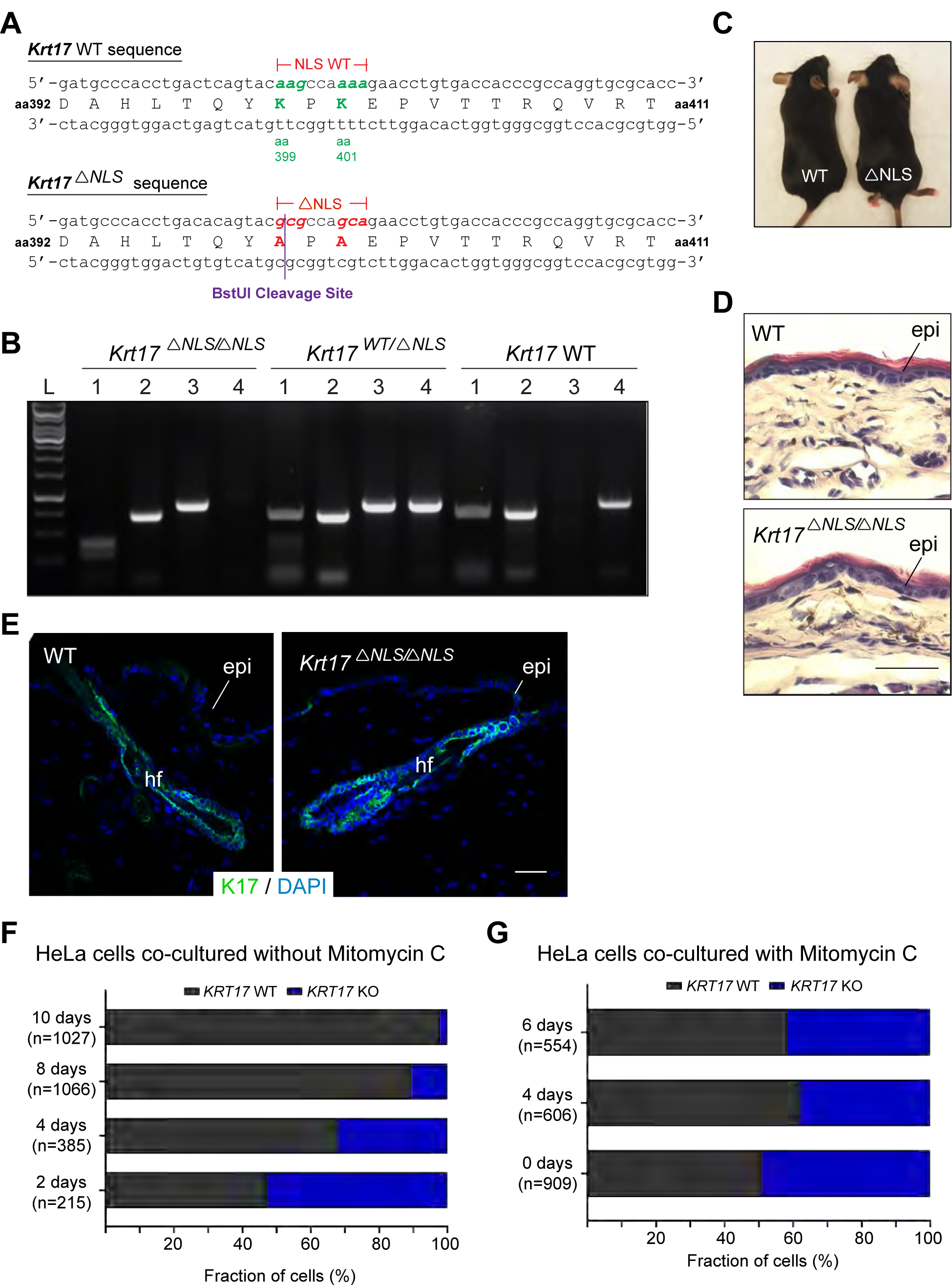
Genesis of a *Krt17*^*ΔNLS/ΔNLS*^ mouse strain and co-culture experiments *KRT17* WT and *KRT17* KO HeLa cells. **(A)** DNA and protein sequences for the *Krt17* WT and *Krt17*^*ΔNLS*^ alleles. Lysine-399 (K399) and Lysine-401 (K401) are both mutated to Alanine residues (K399A and K401A) in the *Krt17*^*ΔNLS*^ mutant allele. The *Krt17*^*ΔNLS*^ mutant allele contains a new BstUI restriction enzyme cleavage site (purple text and line). **(B)** RT-PCR genotyping results for *Krt17*^*ΔNLS/ΔNLS*^, *Krt17*^*+/ΔNLS*^, and *Krt17* WT mice. (lane 1) BstUI digest of the NLS flanking region; (lane 2) undigested control of the product digested in lane 1; (lane 3) PCR amplification using the *Krt17*^*ΔNLS*^ mutant-specific primer; and (lane 4) PCR amplification using the *Krt17* WT-specific primer. **(C)** Age-matched *Krt17* WT and *Krt17*^*ΔNLS/ΔNLS*^ mutant mice demonstrating a comparable phenotype at baseline. **(D)** Hematoxylin and eosin staining of (P60) (Ear) tissue sections from *Krt17* WT and *Krt17*^*ΔNLS/ΔNLS*^ mutant mice. Scale bars, 50 μm. **(E)** Confocal micrographs of hair follicles from *Krt17* WT and *Krt17*^*ΔNLS/ΔNLS*^ mice, immunostained for K17 (green). DAPI, blue. Scale bars, 50 μm. **(F)** Quantitation of the percent of HeLa *KRT17* WT and *KRT17* KO cells after mixing and plating 1:1 and then culturing in normal growth media for different lengths of time. **(G)** Quantitation of the percent of HeLa *KRT17* WT and *KRT17* KO cells after mixing and plating 1:1 and then culturing in normal growth media supplemented with Mitomycin C for different lengths of time.

### Abbreviations

IF: Intermediate Filament
IP: Immunoprecipitation
INM: Inner Nuclear Membrane
K17: Keratin 17
KRT17: Keratin 17
LAP2: Lamina-Associated Polypeptide 2
MEK: Mouse Epidermal Keratinocyte
NLS: Nuclear Localization Signal
PTM: Post-Translational Modification
WT: Wild-Type

## Materials and Methods

### Cell culture

HeLa (cervical, epithelial) and A431 (vulva, epidermoid) keratinocyte cell lines (source: ATCC) were routinely cultured in normal growth media (NGM), which consists of Dulbecco’s Modified Eagle Medium (DMEM; Life Technologies, REF# 11995-065), 10% Fetal Bovine Serum (FBS; Atlanta Biologicals REF# S11150), 100 units/mL Penicillin and 100 μg/mL Streptomycin (P/S; Life Technologies REF# 15140-122), at 37°C and 5% CO_2_. Cells were routinely passaged with 0.05% Trypsin (Corning, REF# 25-052-Cl) when they were ∼75-80% confluent. The protocols for generating HeLa cells null for *KRT17* and confirming the anticipated genome recombination are identical to those previously used to generate A431 cells null for *KRT17* (Hobbs et al., 2015). Prior to use in these studies, all cell lines were treated with a Plasmocin treatment regimen for 2 weeks and confirmed to be mycoplasma free (data not shown). Newborn mouse skin keratinocytes were initiated in primary culture as detailed in Sections 3.2-3.3 in Wang et al. (2016).

For all microscopy experiments, cells were seeded on cover slips (Globe Scientific Inc., REF# 1404-15) that had been previously coated with rat collagen type I (Corning REF# 354236). At the time of harvest, cells were fixed with 4% paraformaldehyde, rinsed with PBS, and mounted on microscopy slides using mounting medium containing DAPI (Duolink REF# 82040-0005). For indirect immunofluorence detection of K17, cells were permeabilized with ice-cold 0.2% TritonX-100 diluted in PBS for 5 minutes on ice prior to antibody incubation as previously described (McGowan et al., 1998).

For the co-culture competition assay, *KRT17* WT and *KRT17* KO cells (10,000 each per cover slip) were cultured for a defined number of days prior to fixation and mounting.

For transient tranfection experiments, GFP, GFP-K17 WT, or GFP-K17ΔNLS cDNA (pEGFP-C3 vector backbone; Addgene REF# 6082-1) were transfected into cells using FuGENE HD Transfection Reagent (Promega REF# E2311) following manufacturer’s recommendations. Cells were cultured on cover slips, as described above, for 1 day prior to transfection and for 2 days following transfection. Microscopy-based analyses and western immunoblotting assays established that the expression levels of the DNA constructs being compared are highly similar if not the same.

### Generation of *Krt17*^*ΔNLS/ΔNLS*^ mice

A mouse strain, *Krt17*^*□NLS/□NLS*^, carrying K399A and K401A point mutations at the *Krt17* locus, was generated using the RNA-guided CRISPR-Cas9 system as described (Wang et al. 2013). A guide RNA (gRNA) (5’-GTACAAGCCAAAAGAACGTA<u>AGG</u>-3’; PAM motif is underlined) was selected based on having a cut site proximal to codon K399 and low predicted off-target recognition (http://crispr.technology) (Jaskula-Ranga et al., 2016), cloned into the pT7gRNA vector using oligonucleotides (Fwd = 5’-CCATCGATAATACGACTCACTATAGTACAAGCCAAAAGAACGTAG-3’;. Rev = 5’-CATGTTCGGTTTTCTTGCATCAAAATCTCGATCTTTATCGTTCAA-3’), and in vitro transcribed and purified (NEB HiScribe T7 High Yield RNA Synthesis Kit) prior to injection. The homology directed repair (HDR) template was purchased as a 160-nt single stranded Ultramer (IDT), and encoded a AAG (Lys) to GCG (Ala) mutation at codon 399 and a AAA (Lys) to GCA (Ala) mutation at codon 401 of the mouse K17 coding sequence (5’-GCATTAGGGGTGAGGAAGGTACTCTCGGATTTCATCTAATCT CTCCAACTTTTTTTTTCCAGCCTGACTCAGTAC<u>GC</u>GCCA<u>GC</u>AGAACGTAAGGATAT TGGTAACTGAGGGCTGGGGTAGAAGGATGCATGTGGCAGGAATCGCCTAGCAGATTGCTAGG-3’; mutation sites are underlined). The gRNA, Cas9 mRNA, and HDR template were co-injected into C57Bl/6 zygotes by the JHU Transgenic Core Facility. Potential transgenic founders were screened using restriction digestion of PCR product extending beyond the repair template oligonucleotide and findings were confirmed by direct DNA sequencing (data not shown). One male homozygote and one male heterozygote founder were selected and independently backcrossed by mating to C57Bl/6 *wildtype* females for three generations to eliminate potential off-target effects. The *Krt17*^*ΔNLS/ΔNLS*^ homozygotes used in this study resulted from het x het breedings.

### Analysis of shape, size, and surface area of cell nuclei

Cells on coverslips were fixed with 4% paraformaldehyde (Electron Microscopy Sciences, REF# 15710) for 10 minutes at room temperature with gentle agitation on a plate rocker and mounted on microscope slides (VWR REF# 48311-703) with mounting medium containing DAPI. Images of nuclei were acquired with a Zeiss LSM700 confocal laser scanning microscope (40X and 63X objectives). Confocal micrographs were processed to maximum intensity projections (MIPs) in order to quantify two-dimensional area measurements of cell nuclei, using the freehand selections tool in the ImageJ processing software (Schneider et al., 2012). Three-dimensional measurements (e.g. volume, sphericity, and surface area) of confocal micrographs were acquired with the Imaris version 8 microscopy image analysis software (Bitplane; http://bitplane.com). Statistical analyses were conducted using the two-tailed unpaired t-test function on the GraphPad Prism 8 software to calculate p-values.

### Whole-cell lysis, SDS-PAGE, and immunoblotting

Whole cell lysates were prepared in urea sample buffer (8 M urea, 1% SDS, 10% glycerol, 60 mM Tris pH 6.8, 0.005% Pyronin Y, and 5% β-mercaptoethanol [β-ME]), homogenized using QIAshredders (QIAGEN REF# 79656), and subjected to Bradford assay (BIO-RAD REF# 500-0006) prior to SDS-PAGE electrophoresis on a 4-15% gradient gel (BIO-RAD REF# 456-1086). 10 µg of protein lysate was loaded per well. Immunoblotting was performed using nitrocellulose membranes (BIO-RAD REF# 162-0115) and a blocking buffer of 5% milk dissolved in either PBS or Tris-buffered saline (TBS).

Primary antibodies utilized for immunoblotting include: rabbit polyclonal anti-K17 (McGowan et al., 1998), mouse monoclonal anti-lamin A/C (Santa Cruz REF# sc-376248), mouse anti-lamin B1/B2 (“2B2” ; generously provided by Dr. Robert Goldman), mouse anti-pan isoform LAP2 (“LAP12” ; generously provided by Dr. Roland Foisner), rabbit polyclonal anti-emerin (Santa Cruz REF# sc-15378), rabbit monoclonal anti-GAPDH (Cell Signaling Technology REF# 5174), mouse monoclonal anti-acetyl-histone H3 (H3K9Ac + H3K14Ac; Santa Cruz REF# sc-518011), mouse monoclonal anti-acetyl-histone H4 (H4S1Ac + H4K5Ac + H4K8Ac + H4K12Ac; Santa Cruz REF# sc-377520), and mouse monoclonal anti-monomethyl-histone H4 (H4K20me1; Santa Cruz REF# sc-134221). Secondary antibodies for immunoblotting include Donkey anti-Mouse IgG and Goat anti-Rabbit IgG (LI-COR REF# 925-32212 and 926-32211). Immunodetection and image capture occurred by infrared imaging (LI-COR Biosciences). Statistical analyses were conducted using the two-tailed unpaired t-test function on the GraphPad Prism 8 software to calculate p-values.

### Indirect immunofluorescence staining for microscopy and quantitative analysis

Cells were fixed and mounted as described above. Primary antibodies utilized for indirect immunofluorescence include: mouse monoclonal anti-lamin A/C (Santa Cruz REF# sc-376248), mouse anti-Lamin B1/B2 (“2B2” ; generously provided by Dr. Robert Goldman), mouse monoclonal anti-pan isoform LAP2 (“LAP12”), rabbit polyclonal anti-LAP2α (“245-3”), mouse monoclonal anti-LAP2β (LAP17”) (all were generously provided by Dr. Roland Foisner), mouse monoclonal anti-emerin (Santa Cruz REF# sc-25284), mouse monoclonal anti-acetyl-histone H3 (H3K9Ac + H3K14Ac; Santa Cruz REF# sc-518011), mouse monoclonal anti-acetyl-histone H4 (H4S1Ac + H4K5Ac + H4K8Ac + H4K12Ac; Santa Cruz REF# sc-377520), mouse monoclonal anti-monomethyl-histone H4 (H4K20me1; Santa Cruz REF# sc-134221), and rabbit polyclonal anti-K17 (McGowan et al., 1998). Species-appropriate AlexaFluor secondary antibodies (488 or 568) were utilized prior to mounting on slides as described above.

For proximity ligation assays (PLA), fixed cells were double immunolabeled with rabbit polyclonal anti-K17 (McGowan et al 1998) and mouse anti-K17 (Santa Cruz REF# sc-393002) prior to incubation with PLA probes (Sigma-Aldrich REF# DUO92002 and DUO92004) for 60 minutes. Before mounting, cells were gently rinsed multiple times with Wash Buffer A (Sigma-Aldrich REF# DUO82049) prior to DNA oligonucleotide incubation for 30 minutes and amplification for for 100 minutes at 37°C (Sigma-Aldrich REF# DUO92007).

### Subcellular fractionation, K17 co-IP, and mass spectrometry-based proteomics

When cells cultured on 15-cm tissue culture dishes reached 80% confluence, they were rinsed with PBS at room temperature, scraped into ice-cold low salt lysis buffer (10 mM Tris pH 7.5, 1 mM MgCl_2_, 10 mM KCl, plus protease inhibitor cocktails (PIC) that include 1 mM PMSF, 1mM Na_3_VO_4_, 1 mM Na_4_P_2_O_7_, 50 mM NaF, 2 µg/mL antipain, 10 µg/mL aprotinin, 10 µg/mL benzamidine, 1 µg/mL leupeptin, 1 µg/mL chymostatin, and 1 µg/mL pepstatin-A), incubated on ice for 30 minutes, and sheared with a 22-gauge needle to release cell nuclei. After pelleting by centrifugation (800 x g, 5 min, 4°C), supernatant was removed and pelleted nuclei were reconstituted in a high salt lysis buffer (40 mM HEPES pH 7.5, 450 mM NaCl, 1 mM EDTA, plus PIC) and twice incubated on ice for 10 min followed by 1 minute vortex prior to ice water bath sonication (8 cycles total; 1 cycle = 30 seconds on, 30 seconds off) and centrifugation (17,000 x g, 10 minutes, 4°C) to pellet insoluble nuclear debris.The resulting supernatant was subjected to pre-clearing of non-specific-binding proteins via incubation with TrueBlot Anti-Rabbit Ig IP Agarose Beads followed by K17 immunoprecipitation (IP), as described previously (Wang et al., 2016). The concentrated eluate was then analyzed by immunoblotting or submitted to the Mass Spectrometry and Proteomics Core Facility at the Johns Hopkins School of Medicine for identification of co-immunoprecipitated proteins.

### Mass spectrometry and bioinformatics analysis

For mass spectrometry (MS), the immunoprecipitation eluates were each adjusted to pH 8.0 using 500 mM TEAB buffer, reduced with 16 mg/mL DTT for 1 hour at 60°C, and alkylated with 36mg/mL Iodoacetomide for 15 minutes at room temperature under low-light conditions. Proteins were then precipitated by TCA/acetone extraction overnight at - 20°C, reconstituted in 100 mM TEAB, and proteolyzed into peptide fragments with 1 µg of lyophilized trypsin overnight at 37°C. Tryptic peptides were then labeled with TMT 10pplex isobaric tags (ThermoFisher REF# 90110; TMT10-127N and TMT10-130N reagents were used to label the eluates from the *KRT17* WT and *KRT17* KO cell lysates, respectively) for 1 hour at room temperature. TMT-labeled peptides were then mixed together and step-fractionated by basic reverse-phase (bRP) chromatography (Waters Oasis u-HLB plates) using 10 mM TEAB containing 5%, 15%, 20%, 30%, and 75% acetonitrile. The resulting 5 bRP fractions were evaporated until dry and then reconstituted in 2% acetonitrile/0.1% formic acid. Finally, samples were analyzed by liquid chromatography/tandem mass spectrometry (LC-MS/MS). Labeled peptides were first separated by a nano-Acquity LC system (Waters) using reverse-phase chromatography on a 75-um x 150 mm ProntoSIL-120-5-C18 H column (2%-90% acetonitrile/0.1% formic acid gradient over 90 minutes at 300 nL/min) and then detected using a Q Exactive HF in FTNT Quadrupole-Orbitrap mass spectrometer (Thermo Fisher Scientific). Tandem MS/MS spectra were processed by Proteome Discoverer software (ThermoFisher Scientific, v1.4) and analyzed with Mascot v.2.5.1 Matrix Science (www.matrixscience.com) using 2015RefSeq_72r_human_plus (human peptide database).

The raw MS data was subject to appropriate filtering prior to analysis, as follows. Entries with a 127N/130N isobaric tag labeling ratio equal to or greater than 1.20 (n = 140), which reflects enrichment over negative control K17 IP from *KRT17* KO cells, along with a 127N/130N isobaric tag count equal to or greater than 4 (n = 77), were selected for further analysis. Of the top 77 proteins that persisted through this filtering, 58 (75%) had at least one defined nuclear role according to Gene Ontology terms in the Universal Protein Resource (UniProt) database (https://www.uniprot.org/). These 77 proteins were then subjected to association network analysis using the online STRING-db freeware (Jensen et al., 2009, Szklarczyk et al., 2015).

### Luciferase assays

Renilla luciferase control plasmid pRL-TK (Promega, E2241), GLi activity responsive Firefly luciferase plasmid 8xGLiBs-luc (Bianchi et al., 2005), the pEGFP-C3 K17 WT (Pan et al., 2011), the pCIG empty vector (Holtz, 2013), the GLi1-pCIG over-expression vector (Carpenter, 2015), and the GFP-K17ΔNLS (Hobbs et al., 2015) were nucleofected into HeLa cells using SE Cell Line 4D X nucleofector Kit S (Lonza #V4XC-1032) with setting DS-138, plated in a 96 well, and subjected to the Promega Dual Luciferase Reporter Assay System (Promega, PR-E1910) to measure Firefly and Renilla luciferase activities 24 hours post nucleofection. Firefly relative light unit (RLU) was normalized to internal Renilla RLU per well after measurements were recorded. Five biological replicates of normalized Firefly RLUs were combined for each parameter, and the means of each parameter were compared using a Mann-Whitney test. Data presented were transformed by dividing individual RLUs of each parameter by the mean of pRL-TK control parameter alone which was then subjected to statistical analysis.

## Author Contributions

Justin T. Jacob conducted most of the experiments and subsequent data analysis and quantitation. Brian G. Poll generated the knock-in mouse and conducted the validation and initial characterization. Raji R. Nair isolated, cultured, and immunostained primary mouse epidermal keratinocytes and conducted the RT-qPCR analysis of GLI1 target genes. Ryan P. Hobbs, Michael J. Matunis, and Pierre A. Coulombe assisted in study design and data interpretation. Christopher Piñeda performed the luciferase assays. Justin T. Jacob and Pierre A. Coulombe jointly wrote the manuscript.

